# SraTailor: GUI software for visualizing high-throughput sequence read archives

**DOI:** 10.1101/005231

**Authors:** Shinya Oki, Kazumitsu Maehara, Yasuyuki Ohkawa, Chikara Meno

## Abstract

Raw high-throughput sequence data are deposited in public databases as SRAs (Sequence Read Archives) and are publically available to every researcher. However, in order to graphically visualize the sequence data of interest, the corresponding SRAs must be downloaded and converted into BigWig format through complicated command-line processing. This task requires users to possess skill with script languages and sequence data processing, a requirement that prevents a wide range of biologists from exploiting SRAs. To address these challenges, we developed SraTailor, a GUI (Graphical User Interface) software package that automatically converts an SRA into a BigWig-formatted file. Simplicity of use is one of the most notable features of SraTailor: entering an accession number of an SRA and clicking the mouse are the only steps required in order to obtain BigWig-formatted files and to graphically visualize the extents of reads at given loci. SraTailor is also able to make peak calls and files of other formats, and the software also accepts various command-line-like options. Therefore, this software makes SRAs fully exploitable by a wide range of biologists. SraTailor is freely available at http://www.dev.med.kyushu-u.ac.jp/sra_tailor/.

## Introduction

Chromatin immunoprecipitation combined with massively parallel DNA sequencing (ChIP-seq) is a powerful method for comprehensively characterizing the occupancy of genomic regions by transcription factors, the transcriptional machinery, modified histones, and other proteins or protein complexes (Furey 2012). ChIP-seq results are often visualized as histograms, with the horizontal axis showing the genomic position and the vertical axis showing the extent of immunoprecipitated and mapped reads (so-called coverage scores; e.g., Fig. S1 blue in Supporting Information). Although many ChIP-seq studies have been published, the figures they contain often show only a limited subset of the information contained in huge datasets. If researchers wish to learn the protein occupancy in any region of interest, they have three choices. One of the easiest ways is to download files describing coverage scores throughout the entire genome. For example, using Wiggle- and BedGraph-formatted text files and BigWig formatted binarized file, coverage scores can be visualized on genome browsers (e.g., Fig. S1 blue in Supporting Information). However, such files are rarely available as supplementary materials attached to papers, due to their huge size (several tens to hundreds of megabytes). The second way is to download peak-call data files, which are available as supplementary materials more often than coverage score files. Peak-call data files are tab-delimited text files in BED format describing genomic regions occupied by a protein in a statistically significant manner, and they can be visualized on genome browsers (e.g., Fig. S1 red in Supporting Information). However, it should be noted that more than ten peak-call algorithms are in common use, and each of these algorithms offers multiple optional parameter settings. Consequently, the resultant peaks are largely dependent on the authors’ choice of peak-call algorithms and the set values of the associated parameters, as well as on the mapping algorithms. An additional concern is that such peak-call data are not necessarily provided in papers. The final way, which is the most difficult but also the most certain to be effective, is to download and process the raw ChIP-seq data. Most publishers require authors to deposit raw sequence data (Sequence Read Archives, SRAs) into public databases such as GEO of NCBI (http://www.ncbi.nlm.nih.gov/gds). Thus, in principle, every researcher can obtain the full dataset of any published ChIP-seq study, and analyze this data to learn the protein occupancy in any region of their interest. To do so, however, interested parties must process SRAs through computational pipelines, programmed in script languages, that entail several steps including de-archiving, mapping, binarization, and peak calling. This requirement represents a significant technical hurdle for researchers who are not intimately familiar with the command-line interface (CLI). Galaxy (Giardine *et al.* 2005; Goecks *et al.* 2010) is a web-based GUI platform that is capable of processing sequence data using multiple tools. Although CLI operation is not demanded in Galaxy, users still must appropriately select, order, and combine these tools in order to perform accurate processing; therefore, use of the platform requires a sufficiently advanced knowledge of sequence data processing.

In order to easily obtain the coverage scores, we developed a GUI software package, SraTailor. The usage of this software does not require any knowledge of sequence data processing: a BigWig-formatted file is obtained simply by entering the SRA accession number of interest, and then clicking the mouse. In addition, SraTailor can handle files in other formats (eg. BED-formatted peak calls and BAM files) and allows various optional settings when operated in CLI mode.

## Results and Discussion

### Overview and basic usage of SraTailor

We developed a Mac Automator application, SraTailor, capable of automatic conversion of SRAs into BigWig files with complete graphical user interface (GUI) operation. At the first execution of SraTailor, free bioinformatics CLI tools are automatically downloaded and installed after the user agrees to each license (shown in blue letters in Figs. 1B and 2 right). After the initial settings are established, execution of SraTailor opens a menu window (Fig. S2 in Supporting Information). At that time, users can automatically download and prepare genomic libraries (whole-genome FASTA, bowtie2 index, and genome-size files) by simply selecting genome assemblies from the list (“Add Genome” in Fig. S2 and Fig. 2). At that point, users are ready to obtain a BigWig file by entering three minimal inputs: a GEO accession number beginning with GSM, a genome assembly onto which the results will be mapped, and arbitrary file name (“Make BigWig” in Fig. S2 and Fig. 1A). The user’s computer then downloads SRAs corresponding to the GSM accession number and converts them into a BigWig file through the command pipelines illustrated in Fig. 1B. The resultant BigWig file can be viewed with a genome browser IGV (Robinson *et al.* 2011; automatically installed at the initial execution) by selecting the “Viewer” button from main menu panel (Fig. S2 in Supporting Information). Coverage scores are expressed as RPM (Reads Per Million mapped reads) unit in BigWig and BedGraph, such that the number of mapped reads on a given position was scaled (i.e. normalized) against total mapped reads (as shown in Fig. S1 blue in Supporting Information). The scaled coverage scores enable users to compare multiple tracks using a common base. Besides NCBI GEO accession numbers beginning with GSM, archives deposited in other databases are also acceptable as input (e.g., NCBI SRA, EBI ENA, and DDBJ DRA accession numbers beginning with SRX, ERX and DRX, respectively).

**Figure 1.**
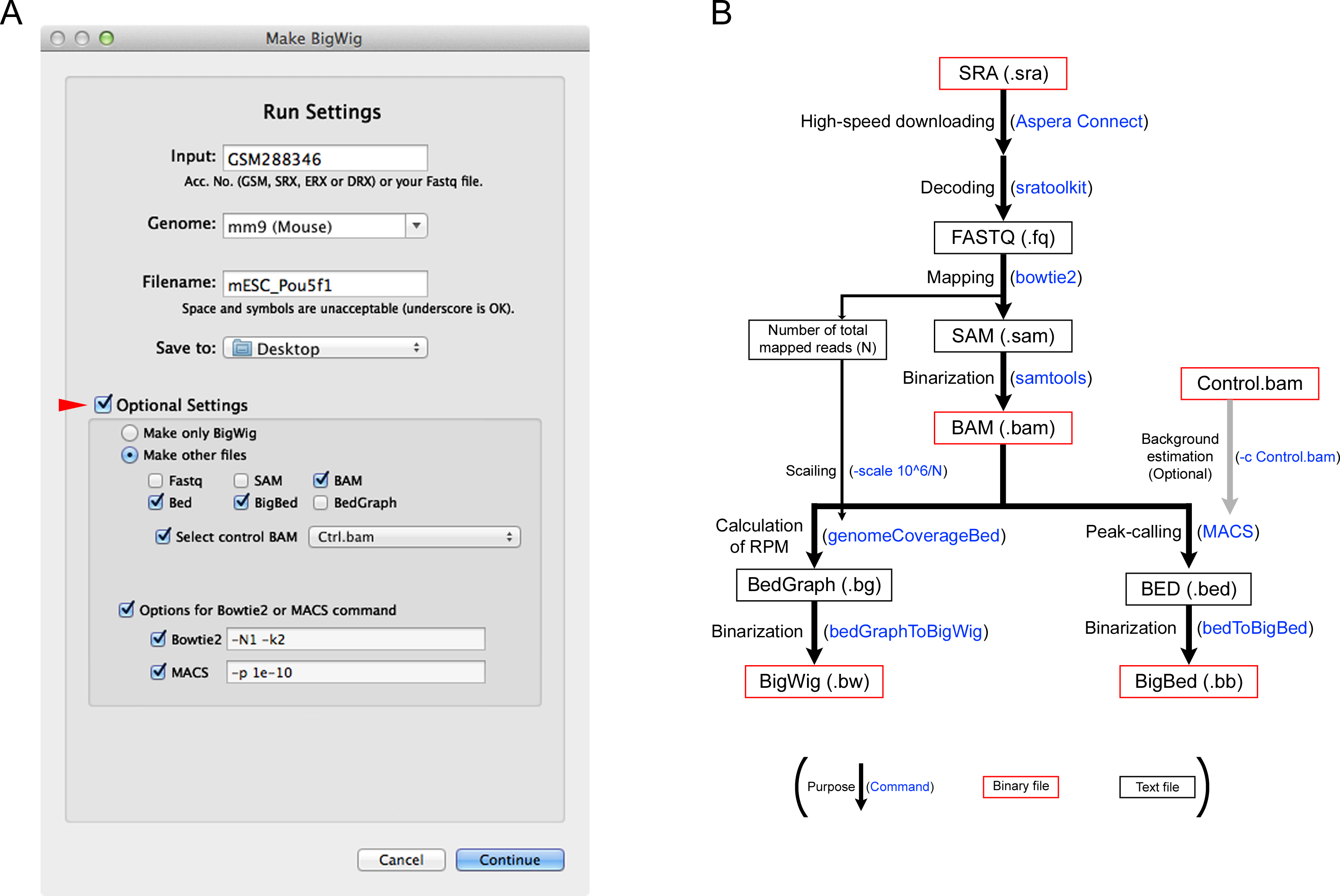
SraTailor “Make BigWig” window. (A) BigWig file is obtained by entering a GEO accession number beginning with GSM (Input), a genome assembly (Genome), and a file name (Filename). Checking “Optional Settings” (arrowhead) opens an additional wizard below, allowing the user to adjust optional settings. (B) After completing the settings, SRAs corresponding to the GSM number are automatically downloaded, followed by conversion into user-requested files through the computational pipelines, as shown. The set of brackets at the bottom provides a legend for the flowchart: black text indicates the purpose of an operation, blue text indicates the command, black text in red box indicates a binary file, and black text in black box indicates a text file. Algorithmic details are described in Materials and Methods.

**Figure 2.**
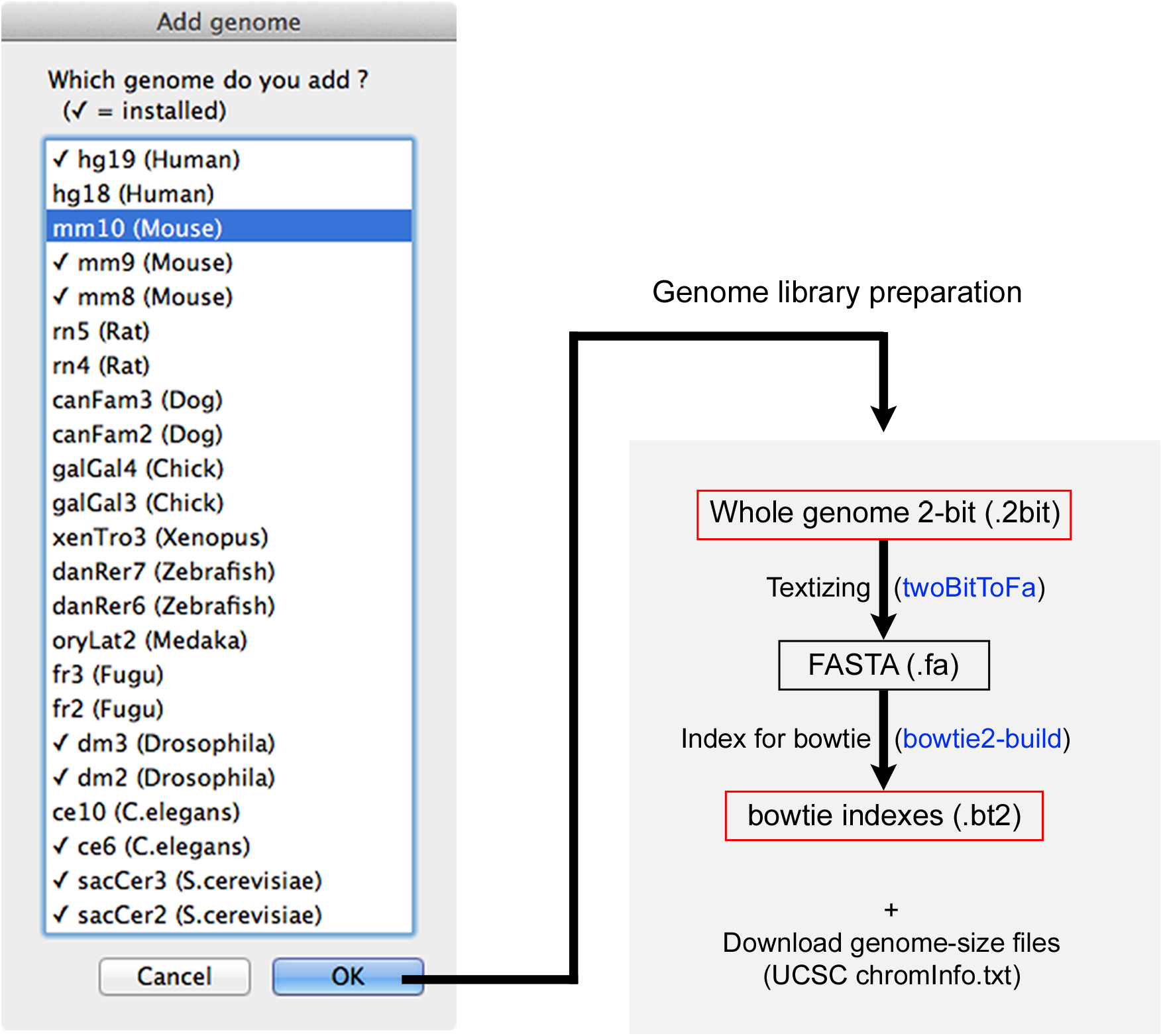
SraTailor “Add genome” window. Genome library files are prepared by selecting the genome assembly from the list. The flowchart represents a method for preparing library files of genome assemblies, according to the set of brackets at the bottom of Fig. 1B. Details of the algorithms are described in Materials and Methods.

### Benchmark test

Benchmark tests for SraTailor were performed on two Macs: iMac (model 13,1) and MacBook Air (model 6,1). We measured the run times required to produce BigWig files from specified GSM accession numbers for mouse EP300 ChIP-seq data from various tissues. Consequently, the approximate computation times were estimated to be 9 and 13 minutes per 10 million reads on iMac and MacBook Air, respectively (Fig. 3). Therefore, SraTailor is able to produce BigWig files within a practical time period on a conventional Mac. More than half of the computation time is spent on mapping with Bowtie2 and conversion to BAM format with SAMTools; both tools can process in multi-thread mode (option `-p` and -@ <thread number>, respectively). SraTailor automatically maximizes the thread number, depending on the Mac on which it is running. Accordingly, the performance depends not only on CPU speed and RAM size but also on the number of CPU cores. The Mac system requirements for SraTailor are described in Doc S1 in Supporting Information.

**Figure 3.**
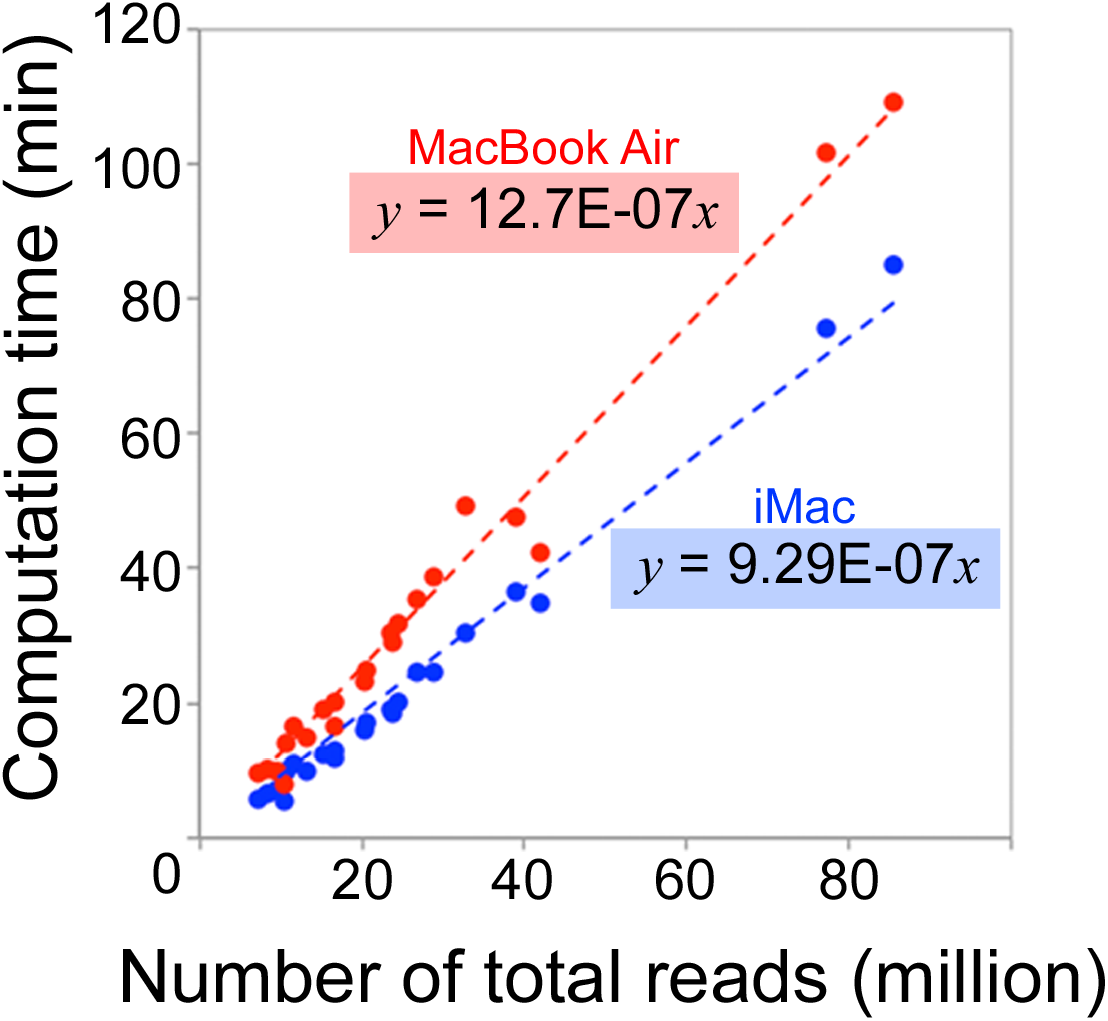
Benchmark test of SraTailor. Run times for producing mm9 BigWig file without “Optional Settings” were measured using an iMac (model 13,1; blue) and a MacBook Air (model 6,1; red). GSM numbers for mouse EP300 ChIP-seq studies (described in Materials and Methods) were tested.

### Advanced usage of SraTailor

As the default, only the BigWig file is produced, but optional settings are also available by clicking the “Optional Settings” check box (Fig. 1A, red arrowhead). For example, users can obtain BigWig and other file formats, such as BAM (in order to obtain the read details; e.g., Fig. S1 gray in supporting information) and BED- or BigBed-formatted peaks (to obtain information about statistical significance). Background estimation is optionally available for peak detection by selecting a pre-made BAM file originating from reads of control input DNA. Customized parameter settings are also possible by entering command-line options for bowtie2 (e.g., `-N1 -k2`) and MACS (e.g., `-p 1e-10`); thus, SraTailor can also satisfy researchers familiar with bioinformatics tools.

In general, high-throughput sequencers offer Fastq files as output, including the nucleotide sequence and data quality. SraTailor also accepts such Fastq files as input, enabling seamless data processing after sequencing. By entering the Fastq file’s path, or mouse-dragging the Fastq file onto the “Input” text field (Fig. 1A), SraTailor can be used to produce files in BigWig and other formats.

SraTailor is principally designed to convert SRAs from high-throughput sequencing into BigWig format. Therefore, it is possible to use SraTailor to visualize coverage of reads originating from RNAs (RNA-seq; excluding the reads at splice junctions), nuclease digests (DNA-seq and MNase-seq), and formaldehyde-unlinked elements (FAIRE-seq) as well as immunoprecipitated fragments (ChIP-seq). Further information on the types of supported datasets is provided in Doc S1 in Supporting Information.

### Concluding remarks

BigWig files are suitable for smooth browsing of coverage scores, and files in this format are now easily obtainable by any researcher via the simple point-and-click interface of SraTailor. While simplicity of operation is one of the most notable features of SraTailor, the software also accepts customized optional settings. SraTailor is a free, open-source application that includes bash script files and Automator workflows, thus allowing anyone to customize the software. SraTailor enables a wide variety of researchers to routinely and fully exploit published ChIP-seq data, and therefore has the potential to accelerate research in the fields of genomics, chromatin, epigenome, and gene-regulatory networks. SraTailor is freely available at http://www.dev.med.kyushu-u.ac.jp/sra_tailor/.

## Experimental procedures

### Software implementation

SraTailor was developed as a Mac Automator application. The contents are shown in Fig. S3 in Supporting Information. Basically, SraTailor is composed of a single Automator “Run Shell Script” action, scripted to execute main.sh file included in SraTailor.app/Contents/sh. The main.sh executes various bash shell scripts (included in SraTailor.app/Contents/sh) and Automator actions and workflows (both included in SraTailor.app/Contents/am) built with Xcode. The shell scripts execute Aspera Connect (http://asperasoft.com) and various CLI tools (included in SraTailor.app/Contents/bin), such as SRA Tool Kit (http://www.ncbi.nlm.nih.gov/sra), Bowtie2 (Langmead & Salzberg 2012), SAMTools (Li *et al.* 2009), BED Tools (Quinlan & Hall 2010), MACS (Zhang *et al.* 2008), and UCSC utilities (http://hgdownload.cse.ucsc.edu/admin/exe/), and they also download and prepare genome library files (included in SraTailor.app/Contents/lib). At the initial launch of SraTailor, initialize.sh is executed in order to download and install the CLI tools mentioned above. Thereafter, launching SraTailor opens a menu panel (Fig. S2 in Supporting Information); clicking on the buttons in this panel results in execution of the following actions:

#### “Make BigWig”

runSettings.sh is executed to open a “Make BigWig” panel (Fig. 1A). The setting parameters are then sent to run.sh, and processing of sequence data is initiated, as shown in Fig. 1B. First, the high-speed downloader Aspera Connect (`ascp` command) downloads SRA files corresponding to the GSM number. The SRA files are then converted into Fastq format using sratoolkit (`fastq-dump` command) and, if the GSM accession includes multiple runs, concatenated into a single file. Reads in the Fastq file are mapped by Bowtie2 in default mode (or with optional user-specified settings), and the resultant SAM file is converted into BAM format using SAMTools (`view` and `sort` command, and optionally `index`). To calculate coverage scores, BedGraph file is produced using Bedtools (`genomeCoverageBed` command) with the `-scale` option in order to express the coverage scores in RPM units. Finally, the BedGraph file is converted into BigWig format using the UCSC utility (`bedGraphToBigWig` command). If peak calls are requested, the BAM file is processed through MACS (`macs14` command) with `-g <genome size>` in default mode (or with optional user-specified settings) in order to produce two BED4-formatted files (peaks.bed and summits.bed) describing genomic positions and MACS peak values. The BigBed files are generated from the BED files using the UCSC utility (`bedToBigBed` command).

#### “Viewer (IGV)”

runIGV.sh is executed to open the IGV genome browser.

#### “Add Genome”

A list dialog is opened (Fig. 2 left), and the returned value is sent to genomeSettings.sh in order to prepare the library files (Fasta-formatted whole-genome sequences, genome sizes, and bowtie2 indexes) as illustrated in Fig. 2 (right). To make a whole-genome FASTA file, a .2bit file of the selected genome is automatically downloaded from the UCSC FTP server (ftp://hgdownload.cse.ucsc.edu) and converted into FASTA format with the UCSC utility (`twoBitToFa` command). The resultant FASTA file is used for producing bowtie2 indexes with the `bowtie2-build` command. Files describing the genome sizes are also downloaded from UCSC FTP server. In other genome assemblies (not illustrated in Fig. 2), FASTA files are generated from archives (downloaded from the UCSC FTP server) using conventional decompression commands (`tar, unzip`, or `gunzip`); alternatively (or in addition), pre-built bowtie2 indexes are directly downloaded from a Bowtie2 index FTP server (ftp://ftp.ccb.jhu.edu/pub/data/bowtie2_indexes/) if available.

#### “Help me!”

A PDF-formatted instruction manual (help.pdf) is opened.

### Benchmark test

Benchmark tests for SraTailor were performed on two Macs: iMac (model 13,1; 2.7 GHz Intel Core i5 with 8 GB RAM) and MacBook Air (model 6,1; 1.3 GHz Intel Core i5 with 4 GB RAM). The run times required to produce BigWig files were measured with following GEO accession numbers as input: GSM1052708, GSM427087, GSM348066, GSM559652, GSM348064, GSM348065, GSM722701, GSM594600, GSM1199037, GSM559653, GSM722762, GSM722862, GSM921138, GSM921137, GSM921134, GSM921136, GSM921135, GSM921139, GSM918750, GSM918747, GSM912893 and GSM912920.

## Acknowledgements

We thank K. Takaoka for testing the software. This work was supported by JSPS KAKENHI Grants 23770254, 25840087, 21116002, and 25670491, and by Kyushu University Interdisciplinary Programs in Education and Projects in Research Development (P&P).

## Supporting Information

**Figure S1** Visualization of BigWig, BED and BAM files on genome browser. Pou5f1 ChIP-seq data in mouse embryonic stem cells (GSM566277; Ang *et al.* 2011) are visualized in the vicinity of *Pou5f1* locus with the genome browser IGV.

BigWig-formatted coverage scores (blue), BED-formatted peak calls (red) and BAM-formatted read details (gray) are shown.

**Figure S2** SraTailor “menu” window. Launching SraTailor opens a menu window. “Make BigWig” button opens the window shown in Fig. 1A, “Viewer” launches the genome browser IGV, “Add Genome” opens a list dialog (Fig. 2) for selecting a genome assembly to be added, and “Help me!” opens a help window.

**Figure S3** File composition of SraTailor.app. Files included in /Applications/SraTailor.app/Contents/ are shown.

**Doc S1** System requirements and supported dataset types

## References

Ang, Y.-S., Tsai, S.-Y., Lee, D.-F. et al. (2011) Wdr5 mediates self-renewal and reprogramming via the embryonic stem cell core transcriptional network. Cell 145, 183–197.

Furey, T.S. (2012) ChIP-seq and beyond: new and improved methodologies to detect and characterize protein-DNA interactions. Nat. Rev. Genet. 13, 840–852.

Giardine, B., Riemer, C., Hardison, R.C., Burhans, R., Elnitski, L., Shah, P., Zhang, Y., Blankenberg, D., Albert, I., Taylor, J., Miller, W., Kent, W.J. & Nekrutenko, A. (2005) Galaxy: a platform for interactive large-scale genome analysis. Genome Res. 15, 1451– 1455.

Goecks, J., Nekrutenko, A. & Taylor, J. (2010) Galaxy: a comprehensive approach for supporting accessible, reproducible, and transparent computational research in the life sciences. Genome Biol. 11, R86.

Langmead, B. & Salzberg, S.L. (2012) Fast gapped-read alignment with Bowtie 2. Nat. Methods 9, 357–359.

Li, H., Handsaker, B., Wysoker, A., Fennell, T., Ruan, J., Homer, N., Marth, G., Abecasis, G. & Durbin, R. (2009) The Sequence Alignment/Map format and SAMtools. Bioinformatics 25, 2078–2079.

Quinlan, A.R. & Hall, I.M. (2010) BEDTools: a flexible suite of utilities for comparing genomic features. Bioinformatics 26, 841–842.

Robinson, J.T., Thorvaldsdóttir, H., Winckler, W., Guttman, M., Lander, E.S., Getz, G. & Mesirov, J.P. (2011) Integrative genomics viewer. Nat. Biotechnol. 29, 24–26.

Zhang, Y., Liu, T., Meyer, C.A., Eeckhoute, J., Johnson, D.S., Bernstein, B.E., Nusbaum, C., Myers, R.M., Brown, M., Li, W. & Liu, X.S. (2008) Model-based analysis of ChIP-Seq (MACS). Genome Biol. 9, R137.

